# Experimental validation of the mechanism of stomatal development diversification

**DOI:** 10.1101/2023.03.22.533739

**Authors:** Yuki Doll, Hiroyuki Koga, Hirokazu Tsukaya

**Author notes:** **Author’s E-mail addresses**: Yuki Doll, Hiroyuki Koga;, Hirokazu Tsukaya.

## Abstract

Stomata are the structures responsible for gas exchange in plants. The established framework for stomatal development is based on the model plant Arabidopsis, but diverse patterns of stomatal development have been observed in other plant lineages and species. The molecular mechanisms behind these diversified patterns are still poorly understood. We recently proposed a model for the molecular mechanisms of the diversification of stomatal development based on the genus *Callitriche* (Plantaginaceae), according to which a temporal shift in the expression of key stomatal transcription factors SPEECHLESS and MUTE leads to changes in the behavior of meristemoids (stomatal precursor cells). In the present study, we have tried to validate this model through genetic manipulation experiments on Arabidopsis. By altering the timing of MUTE expression, we successfully generated Arabidopsis plants with early differentiation or prolonged divisions of meristemoids, as predicted by the model. The epidermal morphology of the generated lines resembled that of species with prolonged or no meristemoid divisions. Thus, the evolutionary process was reproduced via shifting MUTE expression. We also observed unexpected phenotypes, which indicated the participation of additional factors in the evolution of the patterns observed in nature. This study provides novel experimental insights into the diversification of meristemoid behaviors.

**Highlight:** Genetic manipulation in Arabidopsis uncovered how changes in temporal gene expression patterns lead to the diversification of stomatal patterns, providing new experimental insights into the evolution of stomatal development.

## Introduction

Stomata are microscopic pores present on the plant epidermis that function as an adjustable gate for gas exchange. Stomata typically comprise two kidney-shaped guard cells (GCs). The process of stomatal development starts with polygonal stomatal precursor cells termed meristemoids, which typically undergo self-renewing asymmetric divisions, known as amplifying divisions. The meristemoids eventually differentiate into round-shaped guard mother cells (GMCs), each of which produces two GCs through symmetric division. Understanding the developmental process of stomata is important for understanding the physio-ecological properties of extant plants (Harrison et al., 2020), the evolutionary history of land plant lineages (Clark et al., 2022), and the general principles of stem cell regulation in multicellular organisms (Han and Torii, 2016).

To date, extensive studies in developmental genetics have been conducted on the model plant Arabidopsis (*Arabidopsis thaliana*) and have identified the key basic helix-loop-helix (bHLH) transcription factors for stomatal development, namely, SPEECHLESS (SPCH), MUTE, and FAMA (Ohashi-Ito and Bergmann, 2006; MacAlister et al., 2007; Pillitteri et al., 2007). In particular, SPCH and MUTE play pivotal roles in fate determination in the early stages of stomatal development (MacAlister et al., 2007; Pillitteri et al., 2007). SPCH is important for the establishment of meristemoids, as well as maintenance of the amplifying divisions of meristemoids. After several rounds of amplifying divisions, the meristemoid division is terminated by MUTE, and the differentiation of meristemoids to GMCs is triggered (MacAlister et al., 2007; Pillitteri et al., 2007).

Detailed analyses in Arabidopsis have revealed genetic networks for stomatal development that control or are controlled by SPCH and MUTE activity (Qi and Torii, 2018; Zoulias et al., 2018; Lee and Bergmann, 2019). For example, EPIDERMAL PATTERNING FACTOR 1 (EPF1) (Hara et al., 2007) and EPIDERMAL PATTERNING FACTOR 2 (EPF2) (Hara et al., 2009; Hunt and Gray, 2009) are secreted cysteine-rich peptides that play a role in fine-tuning stomatal development. EPF2 is expressed in and secreted from meristemoid mother cells (MMCs) and early meristemoids. The secreted EPF2 inhibits neighboring cells from entering the stomatal lineage by triggering the downstream mitogen-activated protein kinase (MAPK) signaling pathway via its receptors ERECTA and TOO MANY MOUTHS (TMM) (Hara et al., 2009; Hunt and Gray, 2009; Lee et al., 2012,2015; Lin et al., 2017). Complementary to EPF2, EPF1 is expressed in later meristemoids, GMCs, and GCs (Hara et al., 2007, 2009). Secreted EPF1 inhibits meristemoid differentiation and ensures stomatal spacing via its receptors ERECTA-LIKE 1 (ERL1) and TMM (Hara et al., 2007; Lee et al., 2012; Lin et al., 2017). Other well-described factors that play a role in fine-tuning stomatal development are BREAKING OF ASYMMETRY IN THE STOMATAL LINEAGE (BASL) and POLAR LOCALIZATION DURING ASYMMETRIC DIVISION AND REDISTRIBUTION (POLAR) (Dong et al., 2009; Pillitteri et al., 2011), which regulate the polarity of meristemoid divisions. BASL and POLAR are expressed in dividing meristemoids, localize in the cell periphery, and act as a scaffold for either MAPK or glycogen synthase kinase 3-like kinase BRASSINOSTEROID INSENSITIVE 2 signaling to regulate SPCH activity (Zhang et al., 2015, 2016; Houbaert et al., 2018). In addition to the upstream factors of SPCH and MUTE, recent studies have also identified their direct downstream targets, which further illuminate the key roles of SPCH and MUTE in orchestrating stomatal development in Arabidopsis (Lau et al., 2014; Han et al., 2018).

The molecular mechanisms of stomatal development have been extensively studied in Arabidopsis, but other types of stomatal ontogenies characterized by different meristemoid behaviors during development and by different patterns of cell arrangement around the mature stomata have also been identified in land plants (Fryns-Claessens and van Cotthem, 1973; Esau, 1977; Payne, 1979; Rudall et al., 2013). In the case of Arabidopsis, meristemoid division exhibits a helical pattern and results in the formation of stomata with three neighboring cells of different sizes (Zhao and Sack, 1999), termed “anisocytic” stomata, which is typical of Brassicaceae species (Pant and Kidwai, 1967). Meristemoids in some taxa, including Begoniaceae and Crassulaceae, undergo further rounds of amplifying divisions in a helical manner that results in stomata surrounded by a helix of four or more cells, termed “helicocytic” stomata (Payne, 1970; Rudall et al., 2017). Some other species, such as those in Ranunculaceae, entirely lack amplifying division, and their meristemoids directly differentiate into GMCs. Consequently, stomata with no special surrounding structures are formed, and these are termed “anomocytic” stomata (Pant and Mehra, 1964). Because the developmental studies at the molecular level are currently confined to a small number of model species, the molecular basis for the diversification of meristemoid behaviors is still poorly understood.

Stomatal key transcription factors SPCH and MUTE are primary candidates for the factors underlying the diversity in meristemoid behavior, since these transcription factors seem to play dominant roles in determining when meristemoid division is initiated and terminated (MacAlister et al., 2007; Pillitteri et al., 2007; Triviño et al., 2013). Some researchers have proposed that divergence in the function or activity of these two transcription factors may drive the differences in stomatal development patterns (Peterson et al., 2010; Rudall et al., 2013). This view is supported by recent findings that SPCH and MUTE orthologs have conserved functions in stomatal development in broad lineages of land plant species (Liu et al., 2009; Danzer et al., 2015; Chater et al., 2016; Raissig et al., 2016, 2017; McKown et al., 2019; Ortega et al., 2019; Wang et al., 2019). In addition, our recent results on the genus *Callitriche* (Plantaginaceae) also indicate the roles of SPCH and MUTE in the stomatal pattern diversification (Doll et al., 2021a) (Fig. 1). *Callitriche* is an ecologically diverse genus comprising not only terrestrial and aquatic species but also amphibious species that can grow both in water and on land (Ito et al., 2017). Accordingly, *Callitriche* species exhibit diversity in various aspects of stomatal development regulation, such as response to submergence (Koga et al., 2020, 2021), dorsiventral stomatal distribution (Doll et al., 2021b), and meristemoid behaviors (Doll et al., 2021a). We found that meristemoids in terrestrial *Callitriche* species underwent multiple rounds of division before differentiating into stomata in a similar way to species with anisocytic stomata. In contrast, meristemoids in amphibious *Callitriche* species skipped the division step and directly formed anomocytic stomata (Fig. 1). This intrageneric diversity in meristemoid behavior was accompanied by differences in the expression patterns of *SPCH* and *MUTE* orthologs in these species. Terrestrial species, which are characterized by amplifying divisions, expressed *SPCH* in the early stage of stomatal development and then expressed *MUTE* after a certain time lag. On the other hand, amphibious species, which did not exhibit amplifying divisions, expressed SPCH and MUTE almost simultaneously with little time lag (Fig. 1). These results were sufficient for constructing a model of the molecular basis for this diversity. The model demonstrated that meristemoids in terrestrial *Callitriche* expressing SPCH can undergo amplifying divisions until the division terminator MUTE is expressed, while those in amphibious species do not divide further because MUTE is immediately expressed, terminates divisions, and induces differentiation into GMCs. Based on these features, we proposed a model of stomatal development diversification in which the evolutionary shift in the temporal pattern of SPCH and/or MUTE expression leads to the loss or gain of meristemoid divisions (Doll et al., 2021a). However, this model lacks sufficient experimental evidence due to difficulties in genetic manipulation of *Callitriche*. More specifically, it is still unclear whether the expression shifts in SPCH or MUTE alone are sufficient to cause an evolutionary change in stomatal patterning.

**Figure 1.**
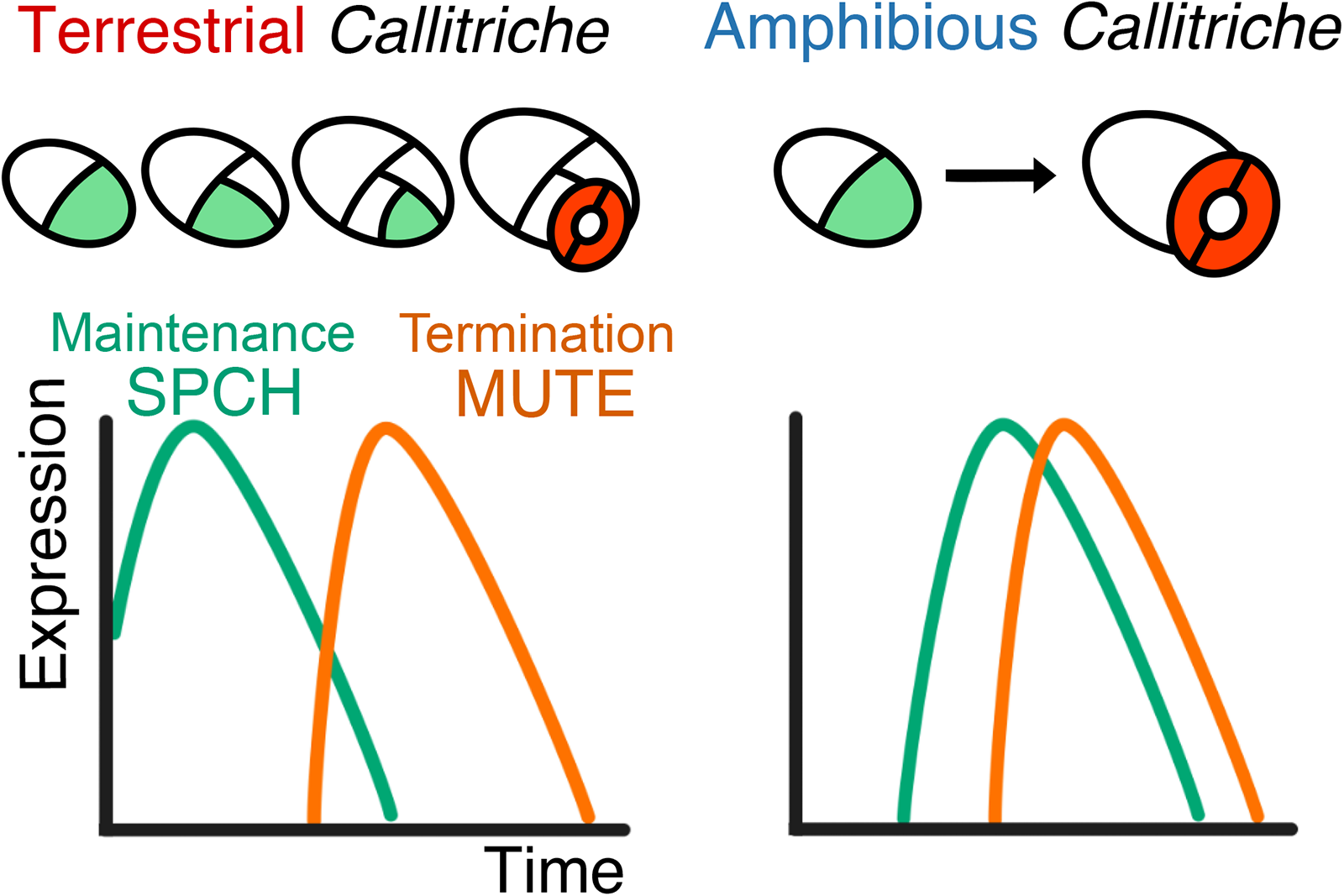
Graphic representation of the hypothetical molecular basis for intrageneric diversity of meristemoid behavior in the genus *Callitriche*. This diagram is reproduced and slightly modified from our previous study, with slight modifications (Doll et al., 2021a). According to this model, differences in meristemoid behaviors between terrestrial and amphibious *Callitriche* species can be attributable to differences in the temporal expression patterns of the key stomatal transcription factors SPCH and MUTE. Meristemoids are indicated in light green, and guard cells are indicated in red.

In the current study, we have tested the validity of the previous model we proposed by manipulating the temporal expression of MUTE in Arabidopsis and examining whether the pattern of stomatal development in this species is modified as predicted by our model. We examined (1) whether induction of early MUTE expression in wild-type (WT) Arabidopsis, characterized by anisocytic stomata, would lead to loss of amplifying divisions (that is, early termination of divisions) and induce transition to anomocytic stomata, and (2) whether delayed MUTE expression would lead to an increase in amplifying divisions (that is, delayed termination of divisions) and induce transition to helicocytic stomata. In order to do so, we established transformant Arabidopsis lines in which MUTE expression is delayed or occurs early (Fig. 2A). By closely examining the phenotypes of the transformants, we assessed the effectiveness and limitations of the model and proposed the mechanisms behind stomatal pattern diversification.

**Figure 2.**
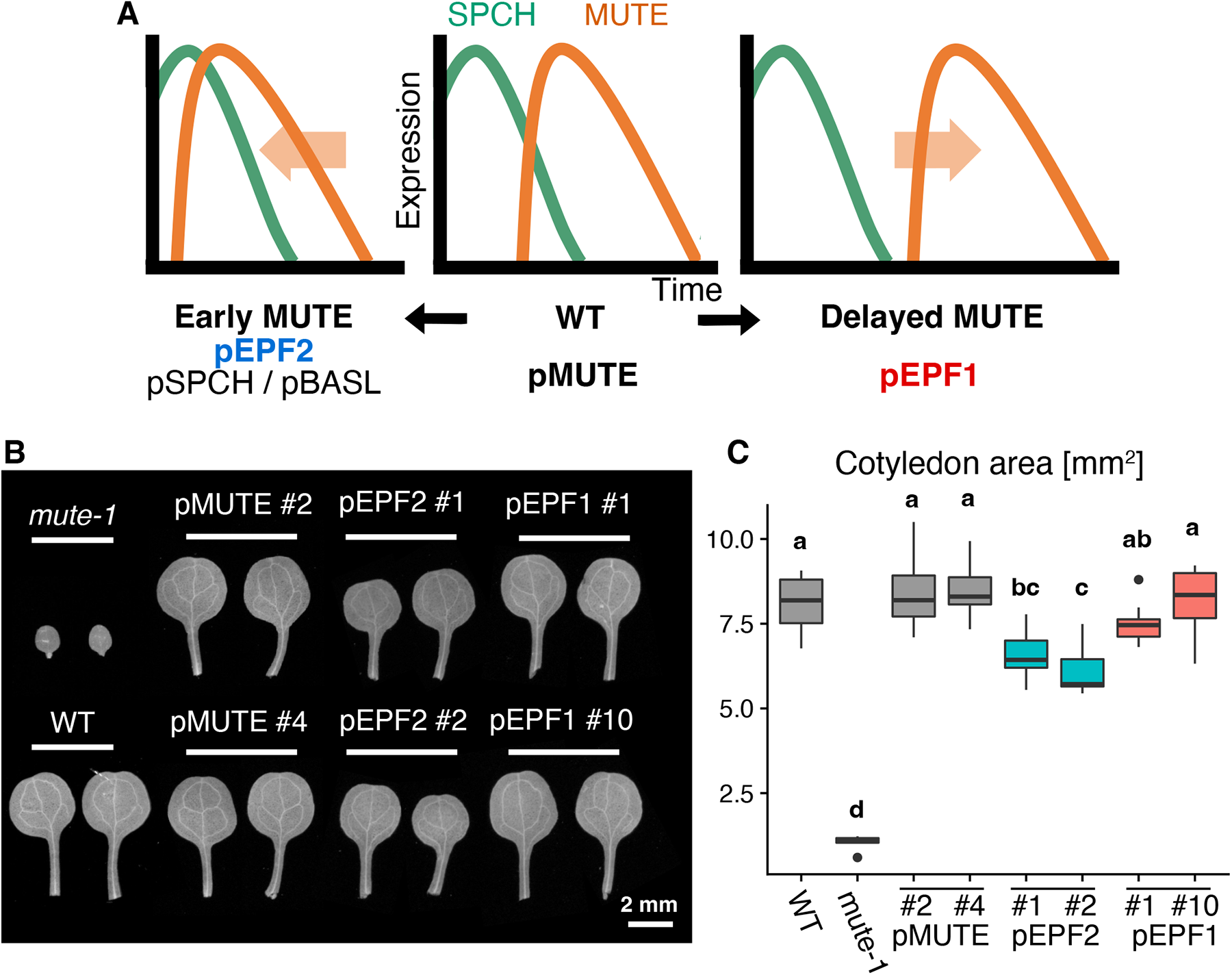
Establishment of the Arabidopsis lines with early or delayed MUTE expression. Schematic view of the experimental design. (**B**) Cotyledon morphology at 20 DAS of each of the established Arabidopsis lines, WT, and *mute-1*. (**C**) Comparison of the cotyledon area at 20 DAS. Groups labeled with different letters are significantly different (p < 0.05) according to the Tukey-Kramer test (n = 10).

## Materials and methods

### Establishment of transgenic Arabidopsis lines

Binary vectors for agrobacterium-mediated transformation were constructed using the backbone of pRI-201 (TaKaRa Bio) or pGWB (Nakagawa et al., 2007) series plasmids. The coding sequence of *AtMUTE* and promoter sequences (*AtBASL*, *AtEPF1*, *AtEPF2*, and *AtSPCH*) were PCR-amplified from Arabidopsis Col-0 cDNA or genomic DNA with the primers listed in Table S1. The promoter of each gene was defined as a 2000–3000 bp long region upstream of the transcription start site, based on previous studies (Dong et al., 2009; Hara et al., 2009; Hara et al., 2007; Pillitteri et al., 2007). The In-Fusion system (TaKaRa Bio) was employed to assemble the amplified sequences and *green fluorescent protein* (*GFP*) sequence with the vector backbones. The resultant vectors and the previously described vector for expression of pMUTE::MUTE-GFP (pLJP155; Pillitteri et al., 2007) were transformed to the *Agrobacterium* C58C1 Rif^r^ strain by electroporation. Transformation to *mute-1* heterozygous Arabidopsis (Pillitteri et al., 2007) was performed with the floral dip method (Clough and Bent, 1998). Stable T4 lines homozygous for *mute-1* and the transgene were used for the following analyses.

### Plant growth and observation of epidermal phenotypes

Transgenic Arabidopsis plants and accession Col-0 WT Arabidopsis were sown and grown on rockwool (Grodan) under 50 μmol m^-2^ s^-1^ white light at 22°C. The surface of the rockwool was covered with the powder of peat moss (Golden Peat Ban; Sakata Seed), and plants were supplied with 0.5 g L^−1^ Hyponex powder fertilizer (Hyponex Japan) in water. To observe mature cotyledons and the first foliage leaves, at 20 days after sowing (DAS), we fixed the plants in the formalin acetic acid-alcohol (FAA) fixative (2.5% formalin, 2.5% acetic acid, and 45% ethanol [v/v]) overnight at 4°C. The fixed mature foliage leaf and cotyledon samples were transferred to an ethanol series (from 70% to 100% [v/v]), treated in thiodiethanol (TDE) solution overnight to clear the tissue (modified from the TOMEI method; Hasegawa et al., 2016), and observed by differential interference contrast microscopy (DM4500, Leica Microsystems). Developing first foliage leaf samples were fixed in FAA solution and treated with Calcofluor White (Sigma-Aldrich) in ClearSee solution (Kurihara et al., 2015) for 2 days to stain cell walls before observation with a confocal microscope (FV10i, Olympus) with UV excitation. Fiji (version 1.0) (Schindelin et al., 2012) was used for analyzing the acquired images.

### Time-lapse observation of epidermal impressions

Time-course series of epidermal impressions of Arabidopsis first foliage leaves were obtained following the dental impression medium-based method described previously (Doll et al., 2021a). To make the molds, we covered the abaxial surface of the leaves with freshly prepared dental impression medium (Provil novo Light, Heraeus Kulzer). After the medium had solidified, it was gently peeled off with fine forceps. In this procedure, molds were made for each leaf on the mornings of the 7th, 8th, 9th, 10th, and 12th day after sowing. Nail polish (AC Quick-Dry Topcoat, Do-Best Inc.) was applied to each mold to obtain a time-course series of the casts. The obtained casts were directly mounted onto glass slides and imaged under a light microscope (DM4500, Leica Microsystems).

### Statistical analysis

All statistical analyses were performed using R (version 4.2.2). The significance level was set at 0.05.

## Results

### Construction of Arabidopsis lines with early or delayed MUTE expression

We induced MUTE expression in the *mute* loss-of-function mutant background (*mute-1*; Pillitteri et al., 2007) by promoters that are active in different stages of stomatal lineage cells, so that MUTE expression is delayed or occurs earlier than that in WT cells (Fig. 2A). To select promoters suitable for this experimental design, we searched previous *in situ* hybridization, promoter assay, and single-cell RNA-seq studies (Liu et al., 2020; Lopez-Anido et al., 2021) for information about the timing of native mRNA expression or promoter activity. We decided to use the *EPF1* promoter (*pEPF1*) to induce delayed MUTE expression, as *pEPF1* is active in late meristemoids and GMCs (Hara et al., 2007). Single-cell RNA-seq analysis has demonstrated that *EPF1* mRNA expression is detected in later stages of the stomatal lineage than *MUTE* mRNA expression (Lopez-Anido et al., 2021). In addition, another single-cell RNA-seq study used *EPF1* as a marker of development stages after the late meristemoid stage (Liu et al., 2020). *EPF1* is unlikely to be under the direct control of SPCH (Lau et al., 2014), but *EPF1* expression is detected even in the *mute* mutant background (Pillitteri et al., 2011). Therefore, we assumed that *pEPF1* could replace the native *MUTE* promoter. For inducing early expression of MUTE, we used the *EPF2 (pEPF2*), *BASL* (*pBASL*), and *SPCH* (*pSPCH*) promoters. Both *pEPF2* (Hara et al., 2009) and *pBASL* (Dong et al., 2009) are active in MMCs and early meristemoids. A previous single-cell RNA-seq result indicated that compared to *MUTE*, both the *pEPF2* and *pBASL* genes are expressed in the earlier stages of stomatal development (Lopez-Anido et al., 2021). Because the two genes are under the direct control of SPCH (Lau et al., 2014), we expected that *pEPF2* and *pBASL* would be suitable for driving MUTE expression immediately after the onset of SPCH expression. In addition, we used *pSPCH* to achieve strict simultaneous expression of SPCH and MUTE.

We designed a genetic construct containing the *MUTE* coding sequence with the *GFP* gene fused to its 3′ end, and its expression was driven by *pEPF1*, *pEPF2*, *pBASL*, or *pSPCH*. This construct was introduced into Arabidopsis with the *mute* mutant background. The resultant transformants are denoted simply as pEPF1, pEPF2, pBASL, and pSPCH henceforth. We also introduced a construct in which the native *MUTE* promoter and gene are fused with *GFP* (Pillitteri et al., 2007) to the *mute* background so that the resultant transformant (pMUTE) can be used as a control line. We confirmed that all the constructs tested rescued the growth retardation phenotype of *mute* to some extent, based on cotyledon morphology (Figs. 2B–C, S1) and shoot appearance (Fig. S2). While the delayed MUTE expression line pEPF1 had as large cotyledons as WT and the control pMUTE line (Fig. 2B–C), the early MUTE expression lines (pEPF2, pBASL, and pSPCH) had retarded growth with smaller cotyledons (Figs. 2B–C, S1). Among the three early MUTE lines, pBASL and pSPCH exhibited more severe growth retardation and unstable phenotypes among independent insertion lines (Fig. S1, S2). Therefore, we mainly focused on pEPF2 in the remaining experiments, as it had a milder and more stable growth phenotype (Figs. 2B–C, S2).

### Confirmation of the experimental design by MUTE-GFP observation

To determine whether the established transgenic lines exhibit the expected MUTE expression patterns, we observed MUTE-GFP expression throughout the leaf development stage in representative lines of each construct (Fig. 3, S3). We observed and counted the number of GFP-positive cells in the abaxial epidermis of developing first foliage leaves in 7–20 DAS plants under a confocal microscope. We found that pEPF2 had a significantly higher number of GFP-expressing cells even in the earliest stage tested (Fig. 3A, G). This indicates that MUTE-GFP is prematurely expressed in this line. In pEPF2, MUTE-GFP was expressed in clusters of cells. This pattern was different from that observed in the control pMUTE line (Fig. 3B, E), which exhibited MUTE-GFP expression in cells with meristemoid- or GMC-like morphology in a dispersed manner. This clustered distribution of MUTE-GFP-expressing cells in pEPF2 persisted even in the later stages (Fig. 3D), and as a result, the GFP-positive cell percentage in this line was high throughout the course of development (Fig. 3G). The other early MUTE lines, namely pSPCH and pBASL, also showed clustered distribution of GFP-positive cells throughout the leaf development stage, as observed in pEPF2 (Fig. S3). In contrast, clustering of GFP-positive cells was not found in pEPF1(Fig. 3C, F). While the morphology and distribution of GFP-positive cells in pEPF1 were similar to those in pMUTE, the number of GFP-positive cells differed in the early and late stages of leaf development. That is, pEPF1 had fewer GFP-positive cells in the early stages (Fig. 3B–C) and more GFP-positive cells in the late stages (Fig. 3E–F) of leaf development than pMUTE. As a result, the peak of MUTE-GFP-positive cell percentage was delayed in pEPF1 (Fig. 3G). In pMUTE, at 11 DAS, most of the remaining meristemoids expressed MUTE-GFP (Fig. 3E); this meant that they would soon differentiate into GMCs. However, at the same stage, pEPF1 had many meristemoids that had not yet expressed GFP (Fig. 3F); this indicated that MUTE-GFP expression was more delayed in this line. To summarize the results, early expression of MUTE in pEPF2 and delayed expression of MUTE in pEPF1 (Fig. 1A) was confirmed based on MUTE-GFP expression, although pEPF2 had an unexpectedly large number of MUTE-positive cells.

**Figure 3.**
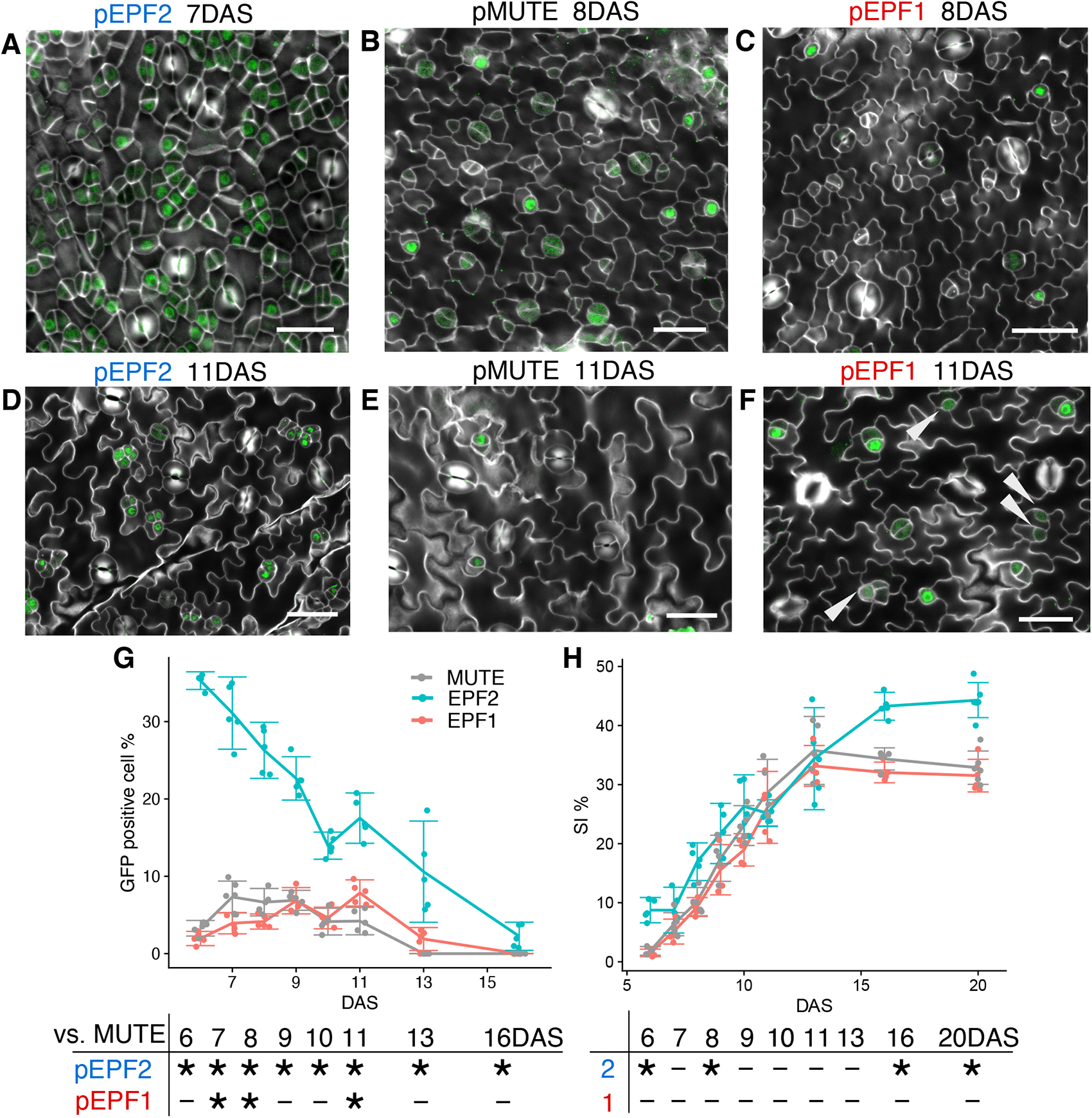
Validation of the experimental design by MUTE-GFP fluorescence observation. (**A–F**) Confocal microscope images of the abaxial surface of developing first foliage leaves in each line. The MUTE-GFP signal is shown in green, and the Calcofluor-stained cell wall is shown in gray. Bars: 25 μm. The images shown are for pEPF2 #1 at 7 DAS (A) and 11 DAS (D), pMUTE #2 at 8 DAS (B) and 11 DAS (E), and pEPF1 #1 at 8 DAS (C) and 11 DAS (F). The white arrowheads in (F) indicate meristemoids without MUTE-GFP signals, which are supposed to be still undergoing amplifying divisions. (**G, H**) Time-course changes in the GFP-positive cell percentage (G) and stomatal index (SI) (H). Each dot represents the data from one leaf primordium. The mean ± 95% confidence interval is shown as an error bar. The table below each graph summarizes the results of the Brunner-Munzel test with Bonferroni correction (*: p < 0.05; n = 5–6) for each time point.

We also measured time-course changes in stomata index (SI), which is defined as the percentage of GCs among all epidermal cells, in order to quantify the number of stomata in each stage of leaf development (Fig. 3H). We found that pEPF2 had significantly higher SI than the control pMUTE line both in the early stages (6 and 8 DAS) and late stages (16 and 20 DAS). The early increase in SI in pEPF2 corresponded to the early peak of MUTE expression in this line. Similarly, the higher SI in the late-stage and mature leaf of pEPF2 reflected a continuously high MUTE-GFP-positive cell percentage in this line. We also found that there was no significant difference in SI between pEPF1 and pMUTE, which implied that comparing the developmental properties of these two lines require closer observation at the cellular level.

### Epidermal morphology of the MUTE-shifted lines resembled helicocytic and anomocytic patterns

After the experimental design was confirmed, we next observed the morphology of the mature epidermis in 20 DAS cotyledons and foliage leaves to determine if the predicted phenotypes of stomatal patterning are shown in each of the transgenic lines (Fig. 4A–C, Fig. S4). In mature cotyledons, consistent with the data obtained for the first foliage leaves (Fig. 3H), SI was significantly higher in pEPF2 than in pMUTE, but there was no significant difference among pEPF1, pMUTE, and WT (Fig. 4D). Notably, pEPF2 had many clusters of stomata, in which each stoma was in direct contact with another stoma (Fig. 4A, E). In WT Arabidopsis and many other plant species, stomata are separated by at least one pavement cell and are rarely arranged adjacent to each other (Sachs, 1991; Geisler et al., 2000), as per the commonly known “one-cell-spacing rule.” Therefore, the clustered stomata in pEPF2 indicate that this line has a defect in the spacing of stomata (which is discussed later). The other early MUTE lines, pSPCH and pBASL, also had clustering stomata, which exhibited a pattern that violated the one-cell-spacing rule (Fig. S4). In contrast, pEPF1 had no clustering stomata (Fig. 4E), which was indicative of appropriate regulation of stomata patterning in this line.

**Figure 4.**
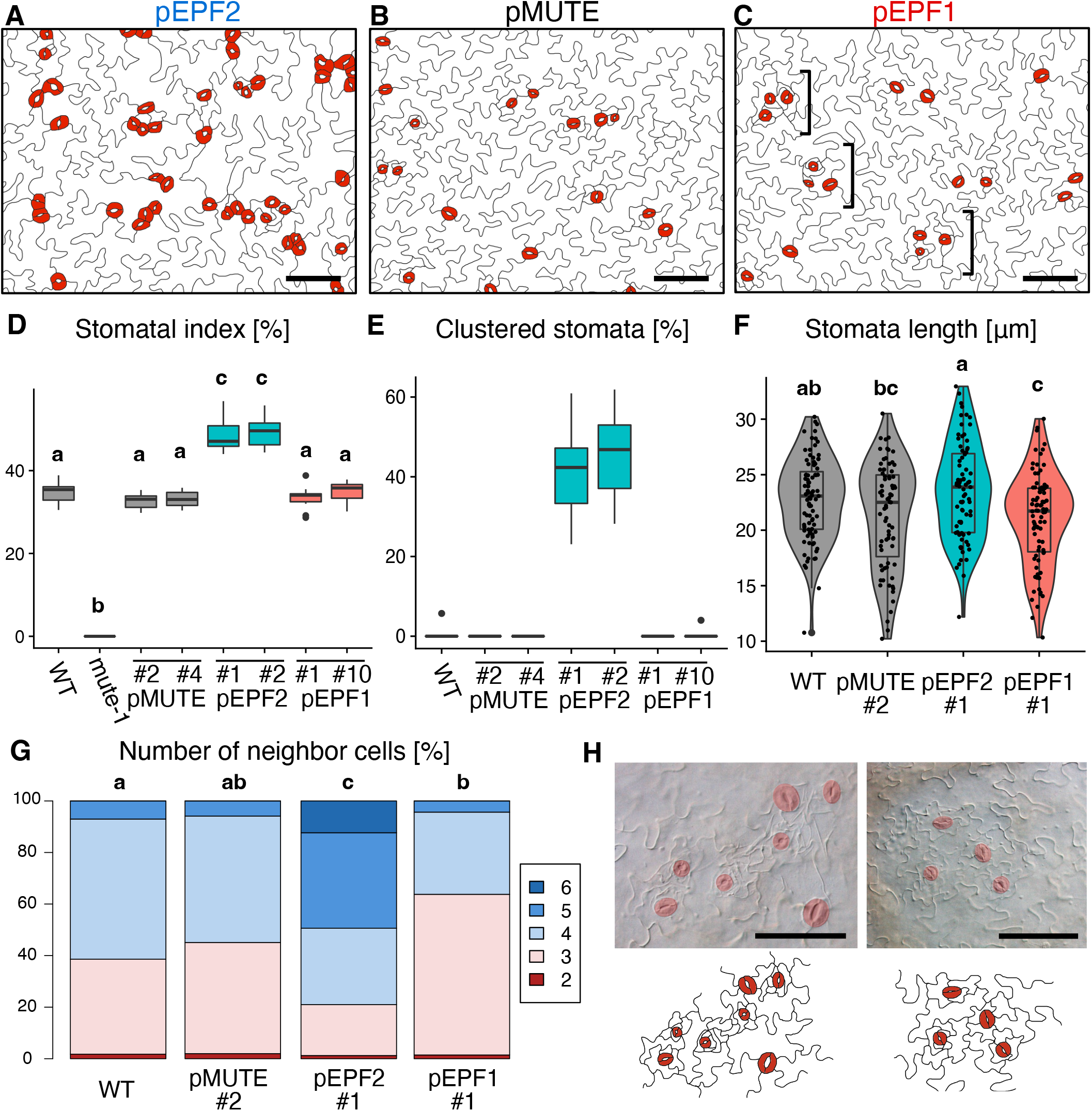
Epidermal morphology of Arabidopsis lines with early or delayed MUTE expression. (**A–C**) Epidermal morphology of mature cotyledons at 20DAS in early MUTE pEPF2 #1 (A), control pMUTE #2 (B), and delayed MUTE pEPF1 #1 (C) lines. Bars: 100 μm. The single square brackets in (C) indicate the clusters of stomata and small pavement cells that resemble non-contiguous stomatal clusters in helicocytic species (see also H). (**D**) Quantification of SI on the abaxial side of 20 DAS cotyledons of the tested lines. Groups labeled with different letters are significantly different (p < 0.05) according to the Tukey-Kramer test (n = 10). (**E**) Quantification of stomata clustering rate on the abaxial side of 20 DAS cotyledons. The number of guard cells in stomata that directly neighbor one or more stomata was divided by the total number of guard cells to calculate the percentage of clustering stomata (n = 10). (**F**) Quantification of stomata size in the abaxial side of 20 DAS cotyledons. Each dot represents data from one stoma. Groups labeled with different letters are significantly different (p < 0.05) according to the Tukey-Kramer test (n > 75 stomata, examined in six different leaves). (**G**) Quantification of the number of cells neighboring stomata on the abaxial side of the 20 DAS first foliage leaves. More than 50 unclustered stomata in three different leaves were examined. Groups labeled with different letters are significantly different (p < 0.05) according to Mood’s median test. (**H**) Examples of the structure that resembles a non-contiguous stomatal cluster in helicocytic species on the abaxial surface of 20 DAS cotyledons of pEPF1 #1. Small pavement cells and stomata are clustered together in accordance with the stomatal one-cell-spacing rule. Bar: 100 μm.

Next, we analyzed morphological traits that may reflect the ontogenic types of the stomata. For these experiments, clustering stomata in pEPF2 (∼40% of all stomata; Fig. 4E) were omitted from the analyses, and only unclustered stomata (∼60%) were examined in order to ignore the effect of defective stomata spacing in this line. We first compared the size of the stomata in terms of length among the tested lines (Fig. 4F). Stomata that differentiated early without division are assumed to be larger than those that have experienced multiple rounds of divisions, since the former would have a longer time for cell expansion, as evidenced by our observation in terrestrial *Callitriche* species showing variation in the size of stomata that differ with regard to the number of amplifying divisions (Doll et al., 2021a). The results showed that pEPF2 had larger stomata, while pEPF1 had smaller stomata, than pMUTE or WT. This implies that pEPF2 may quickly terminate division to start cell differentiation and expansion, while pEPF1 may exhibit delayed onset of cell expansion after many rounds of divisions.

Next, we examined the arrangement of stomata, which reflects the pattern of meristemoid divisions during stomatal development (Fryns-Claessens and Cotthem, 1973; Esau, 1977; Payne, 1979; Rudall et al., 2013) (Fig. 4G). Generally, species with helical meristemoid divisions, including anisocytic and helicocytic species, tend to have stomata with three neighboring cells, whereas anomocytic species without meristemoid division may have stomata with four or more neighboring cells, as demonstrated in the genus *Callitriche* (Doll et al., 2021a). Here, we found that pEPF2 had more stomata with four or more neighboring cells than pMUTE and WT (Fig. 4G). Stomata with three neighbors were less frequent in pEPF2 than in pMUTE and WT. These results indicated that the stomatal arrangement in pEPF2 resembled that of division-less anomocytic stomata. We did not find such a tendency in cluster-forming mutants described in previous studies, when only unclustered stomata were counted (40–85% of all stomata) (Hara et al., 2007; von Groll et al., 2002) (Fig. S5). Therefore, the observed phenotype in pEPF2 is not likely to be the byproduct of stomatal clustering in this line, but rather, it reflects a specific phenotype induced by a temporal change in MUTE expression. Contrary to the observations for pEPF2, in pEPF1, stomata with three neighbors were dominant, with the percentage of such stomata being higher than that in the other lines (Fig. 4G). This indicated that anisocytic or helicocytic stomata arising after helical divisions are dominant in this line. In addition, clusters of small pavement cells and stomata were frequently found in the pEPF1 epidermis (Fig. 4C, H). The distance of each stoma from the nearest neighbor tended to be smaller in pEPF1 compared with WT (Fig. S6). This suggests that stomata in this line are more likely to be present in the proximity of other stomata. Such a cluster is distinct from the stomata cluster found in pEPF2 in that it follows the one-cell-spacing rule. Rather, it is similar to the “non-contiguous stomatal cluster” found in *Begonia* and other helicocytic plant species (Payne, 1970; Rudall et al., 2017).

The morphological observations described above are largely consistent with the prediction from our evolutionary model (Fig. 2A). That is, the delayed MUTE expression line pEPF1 had helicocytic-like traits in terms of both stomatal arrangement and general morphology, and the small size of the stomata was indicative of prolonged divisions (Fig. 4C, F, G, H). In contrast, the early MUTE expression line pEPF2 exhibited phenotypes indicative of early termination of meristemoid divisions, including large stomata and anomocytic-like stomatal arrangement (Fig. 4F, G), though there were also defects in stomatal spacing (Fig. 4A, E).

### Direct observation of meristemoid behavior revealed modified developmental patterns in the MUTE-shifted lines

The above results indirectly support the model being examined. However, different trajectories of stomatal development may produce the same final morphology of mature stomata (Rudall et al., 2013; Rudall and Bateman, 2019). Therefore, direct observation of the developmental process in each transgenic line is necessary to assess the effect of changes in MUTE expression on stomata ontogeny and meristemoid behavior.

To understand the pattern of stomatal development in each line more directly, we observed the behavior of meristemoids in developing leaves by performing dental impression medium-based time-lapse observation (Figs. 5–6). We made serial epidermal impressions of the abaxial side of the first foliage leaves every 1–2 days from 7 DAS to 12 DAS and traced the same epidermal cell population at each time point in order to visualize and quantify the behaviors of each stomatal-lineage cell during this period (Fig. 5). We successfully traced the normal process of meristemoid divisions and its differentiation into a stoma in the control line pMUTE (Fig. 5A, A’). In contrast to the observations in the control line, pEPF2 frequently exhibited abnormal meristemoid behaviors; newly arisen meristemoids and their sister cells (stomatal lineage ground cells [SLGCs]) both differentiated into stomata, forming large contiguous clusters of stomata (Fig. 5B, B’). The divisions producing meristemoid-like cells in pEPF2 were largely asymmetric in appearance (Fig. 5B’), but it is possible that some of them involved abnormally symmetric divisions of meristemoids that produced two neighboring stomatal lineage cells, as observed in division polarity mutants (Dong et al., 2009). Even at 12 DAS, many meristemoids were undifferentiated in pEPF2; this means that abnormal asymmetric or symmetric division of meristemoids lasted over a long period in this line (Fig. 5B). We also detected many meristemoids in pEPF1 at 12 DAS (Fig. 5 C). However, the behavior of meristemoids was normal in this line in that no abnormal stomata differentiation or clustering was found (Fig. 5 C, C’).

**Figure 5.**
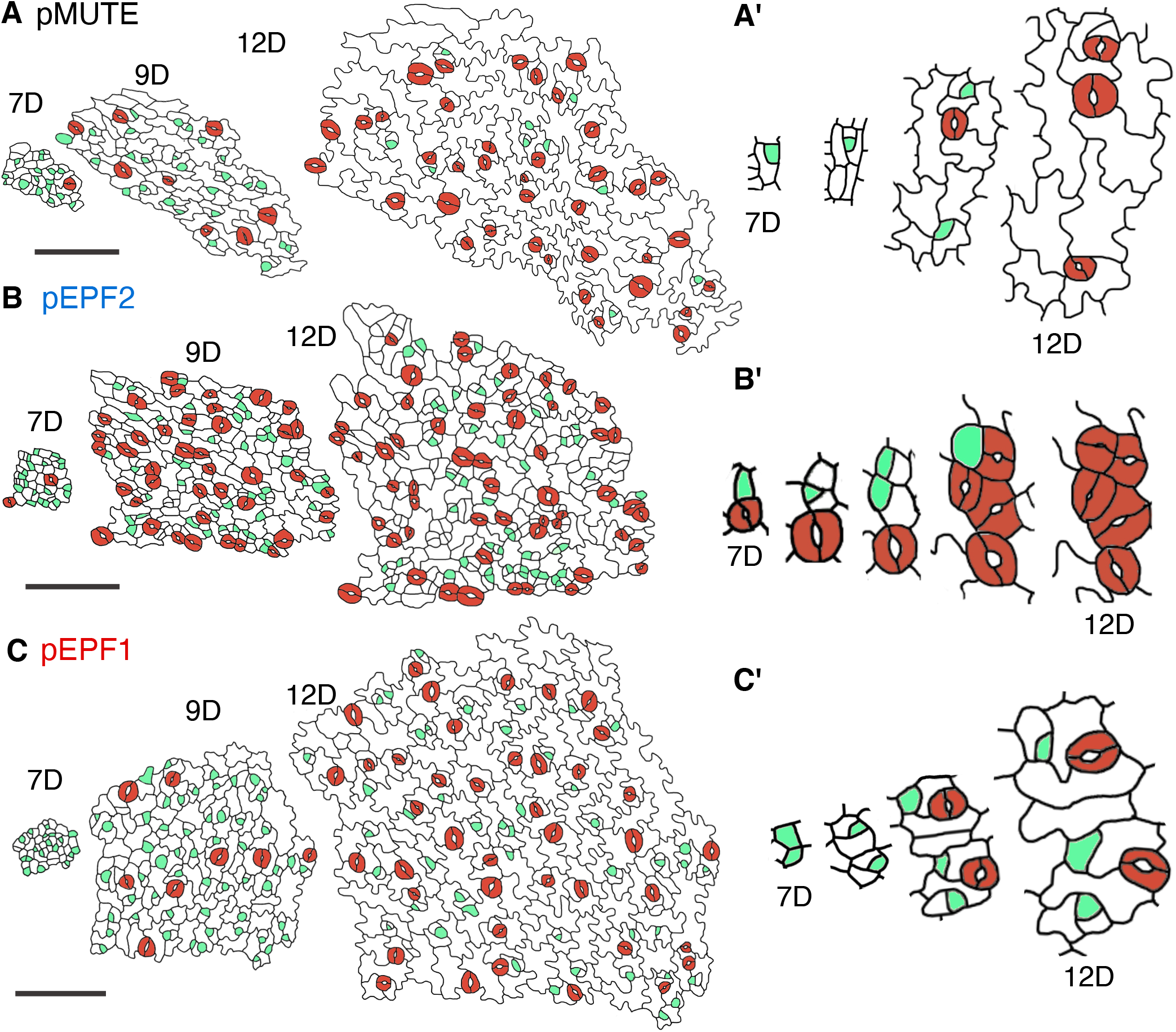
Results of time-lapse analyses of epidermal impressions on the abaxial surface of developing first foliage leaves in Arabidopsis lines with early or delayed MUTE expression. The serial replicas represent impressions from 7 DAS (7D) to 12 DAS (12D). GCs are indicated in red, and stomatal precursor cells, including both meristemoids and GMCs, are in light green. (**A**) Control pMUTE #2 line. (**B**) Early MUTE pEPF2 #1 line. (**C**) Delayed MUTE pEPF1 line. A’–C’ depicts the typical processes of stomatal differentiation in each line, which are taken from a different series of replicas than that shown in A–C. Bars: 100 μm.

**Figure 6.**
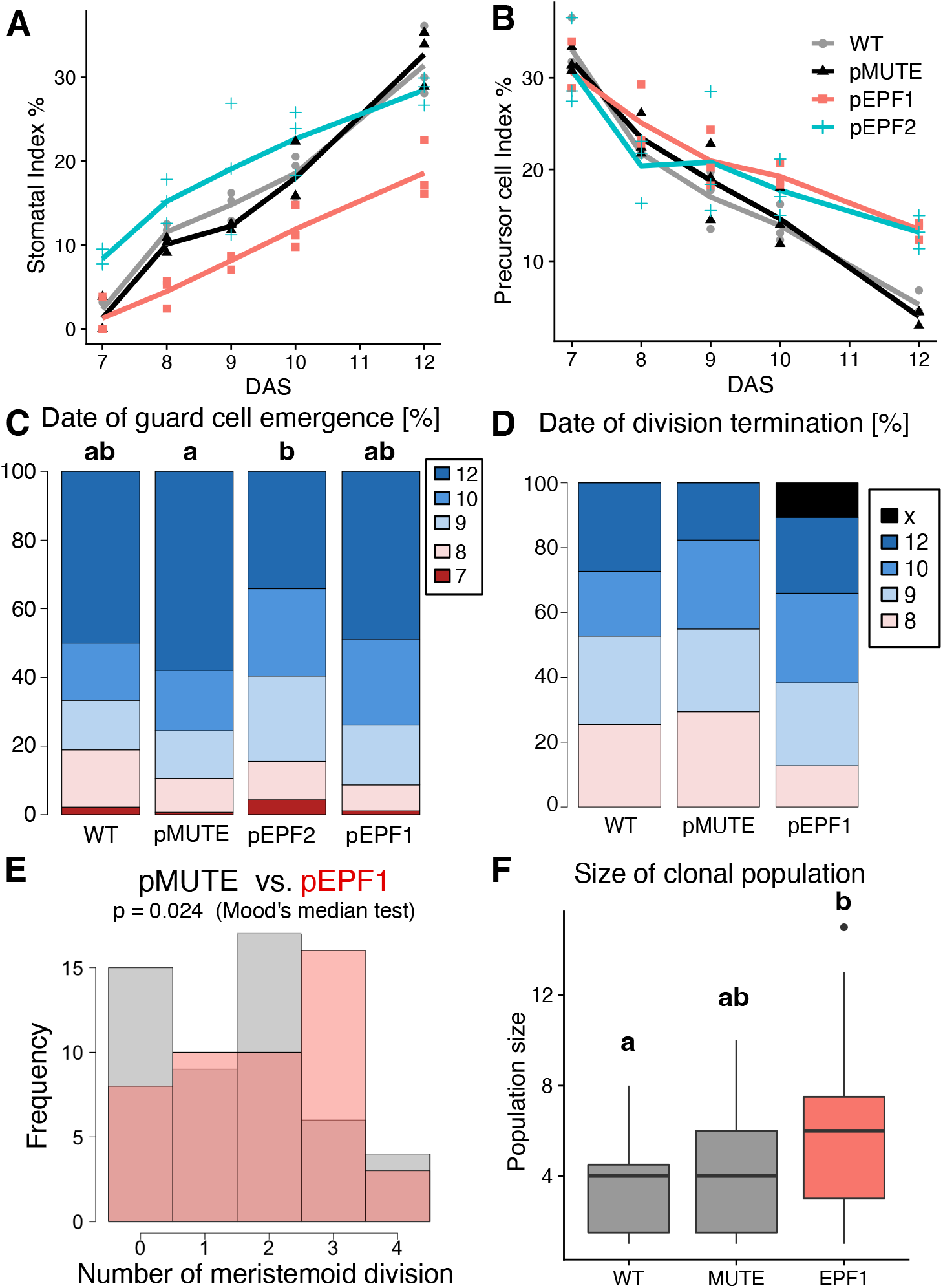
Quantification of time-lapse changes in epidermal impressions of the abaxial surface of developing first foliage leaves in Arabidopsis lines with early or delayed MUTE expression. The representative lines pMUTE #2, pEPF2 #1, and pEPF1 #1 were analyzed as shown in Fig. 5. For each line, three regions from different leaves comprising more than 50 cells at 7 DAS were examined. (**A–B**) Time-course changes in SI (**A**) and PCI (**B**). (**C**) Quantification of the time point of division termination for all the meristemoids identified at 7 DAS (n > 45). The time point at which meristemoids differentiated into GCs was calculated. “x” refers to the meristemoids that were left undifferentiated at 12 DAS. pEPF2 was omitted from the analysis because of the difficulty associated with annotation of meristemoids in this line (see text) (**D**) Comparison of the number of divisions during the observed period (7–12 DAS) for each meristemoid identified at 7 DAS in pMUTE and pEPF1. The p-value according to Mood’s median test (n = 46 for pEPF1 and n = 51 for pMUTE) is shown. (**E**) Comparison of the size of the clonal population that each meristemoid identified at 7 DAS had finally produced by 12 DAS. To minimize the effect of GC production and focus on the division potential of meristemoids, symmetric divisions giving rise to stomata are ignored: the number of GCs is divided by half for calculating the population size (the number of descendant cells for each meristemoid identified at 7 DAS) at 12 DAS. Groups labeled with different letters are significantly different (p < 0.05) according to the Brunner-Munzel test with Bonferroni correction (n > 45). (**F**) Quantification of the time point at which the stomata identified at 12 DAS underwent differentiation. Groups labeled with different letters are significantly different (p < 0.05) according to Mood’s median test (n > 90).

We conducted a quantitative comparison of various parameters related to stomatal development and meristemoid behaviors in each line by analyzing multiple time-course series of epidermal impressions (Fig. 6). We found that pMUTE and WT had almost the same values of SI and precursor cell index (PCI), which refers to the percentage of meristemoids and GMCs among all epidermal cells, throughout the observation period (Fig. 6A–B). This indicates that the developmental pattern of pMUTE is comparable to that of WT. At the start of the observation period (7 DAS), PCI was almost the same for all the lines tested (Fig. 6B). This implies that the leaf developmental stage and the initial patterning of meristemoids (MMCs) in emerging leaf primordia were largely the same among the tested lines.

Quantitative comparison revealed that pEPF2 had larger SI than the other lines in the earliest stages of development (7–9 DAS) compared to the other lines; this is indicative of early differentiation of stomata in pEPF2 (Fig. 6A). This was consistent with the prediction. However, it should be noted that this line had higher PCI at 12 DAS (Fig. 6B). This means that pEPF2 also had active proliferating meristemoids, which may be related to the abnormal behaviors of meristemoids (Fig. 5B, B’). To minimize the effect of such abnormal stomatal differentiation, we focused on unclustered stomata (∼70% of stomata at 12 DAS) in pEPF2 and compared the timing of stomatal differentiation with the other lines. For each unclustered stoma found on the final day of observation (12 DAS), we determined the time point of its emergence (Fig. 6C). The result showed that pEPF2 had more stomata that emerged before 10 DAS than the control pMUTE line. This demonstrated that a portion of the pEPF2 meristemoids differentiated early into stomata without division, as is typically observed in anomocytic species (Doll et al., 2021a; Pant and Mehra, 1964).

The developmental time course of pEPF1 was characterized by low SI throughout the observed period (Fig. 6A) and high PCI at 12DAS (Fig. 6B). Considering that SI in the mature cotyledons and first foliage leaves in pEPF1 was comparable to that in pMUTE and WT (Figs. 3H, 4D), it is assumed that the low SI and high PCI observed in pEPF1 were the results of delayed differentiation of meristemoids. In other words, pEPF1 had many dividing meristemoids left even at 12 DAS, at which point division had been terminated in most meristemoids in WT and pMUTE and they had differentiated into stomata. To determine whether meristemoid division is actually prolonged in pEPF1, we observed meristemoids present at 7 DAS and analyzed their behavior after 8 DAS. pEPF2 was omitted from this analysis because epidermal cells in this line exhibited unusual behavior (Fig. 5B, B’), which made the annotation of meristemoids and non-meristemoid cells at 7 DAS difficult and potentially highly arbitrary. Among the meristemoids found at 7 DAS, all the meristemoids in pMUTE and WT eventually stopped dividing and differentiated into stomata by 12 DAS (Fig. 6D). In contrast, some of the meristemoids in pEPF1 did not stop dividing during the observation period and were still undifferentiated at 12 DAS (Fig. 6D). During the observation period, meristemoids in pMUTE and pEPF1 divided up to four times, with a higher number of divisions observed in pEPF1 than in pMUTE (Fig. 6E). Accordingly, the size of a clonal population at 12 DAS derived from a single meristemoid present at 7 DAS was the largest in pEPF1 among the three analyzed lines (Fig. 6F). These results provide direct evidence that pEPF1, the delayed MUTE line, had a high number of meristemoid amplifying divisions during stomatal development, as is typically observed in helicocytic species (Payne, 1970; Rudall et al., 2017).

## Discussion

In this study, we performed genetic manipulation of Arabidopsis to experimentally validate our model for the evolution of stomatal development that we had previously established based on our analyses in the genus *Callitriche* (Doll et al., 2021a) (Fig. 1). According to this model, changes in the temporal expression patterns of the stomatal key transcription factors SPCH and MUTE, which maintain and terminate meristemoid divisions, respectively, result in an evolutionary change in the behavior of meristemoids. That is, delayed MUTE expression relative to SPCH expression may lead to an increase in meristemoid divisions and the production of helicocytic stomata, while premature expression of MUTE immediately after the onset of SPCH expression may lead to loss of meristemoid division and production of anomocytic stomata. To validate whether such a shift in temporal gene expression patterns can induce changes in stomatal ontogeny, we induced changes in the time point of MUTE expression in Arabidopsis by using different promoters to drive expression of the gene (Fig. 2−3) and analyzed the morphological (Fig. 4) and developmental (Fig. 5–6) phenotypes of the lines. This promoter-based strategy for inducing changes in temporal expression is not suitable for SPCH manipulation, since the identity of the stomatal lineage cell itself is highly dependent on SPCH, making it difficult to select appropriate promoters. Another possible strategy for manipulating temporal expression patterns would be to employ an inducible expression system. For example, a previous study tested the artificial late induction of MUTE using the β-estradiol-inducible expression system (Triviño et al., 2013). However, because the study used the native *MUTE* promoter for driving expression of the estradiol-dependent activator XVE, it was impossible to induce *MUTE* expression earlier than its native expression time point. In addition, due to the intrinsic nature of the inducible systems, long-term and stable control of MUTE expression timings for individual leaves is extremely difficult. Therefore, we used known stomatal lineage-specific promoters for driving MUTE expression at different time points so that the expected temporal pattern of MUTE expression could be stably achieved in individual cotyledons and foliage leaves. Based on our findings for the duration and number of meristemoid divisions, the time point of differentiation into stomatal cells, and stomatal cell arrangement in the pEPF1 line with delayed MUTE expression and the pEPF2 line with early MUTE expression, we can conclude that we were able to successfully modify the stomatal developmental pattern in Arabidopsis. Thus, the proposed model for the diversification mechanism of stomatal development was experimentally validated (Fig. 7).

**Figure 7.**
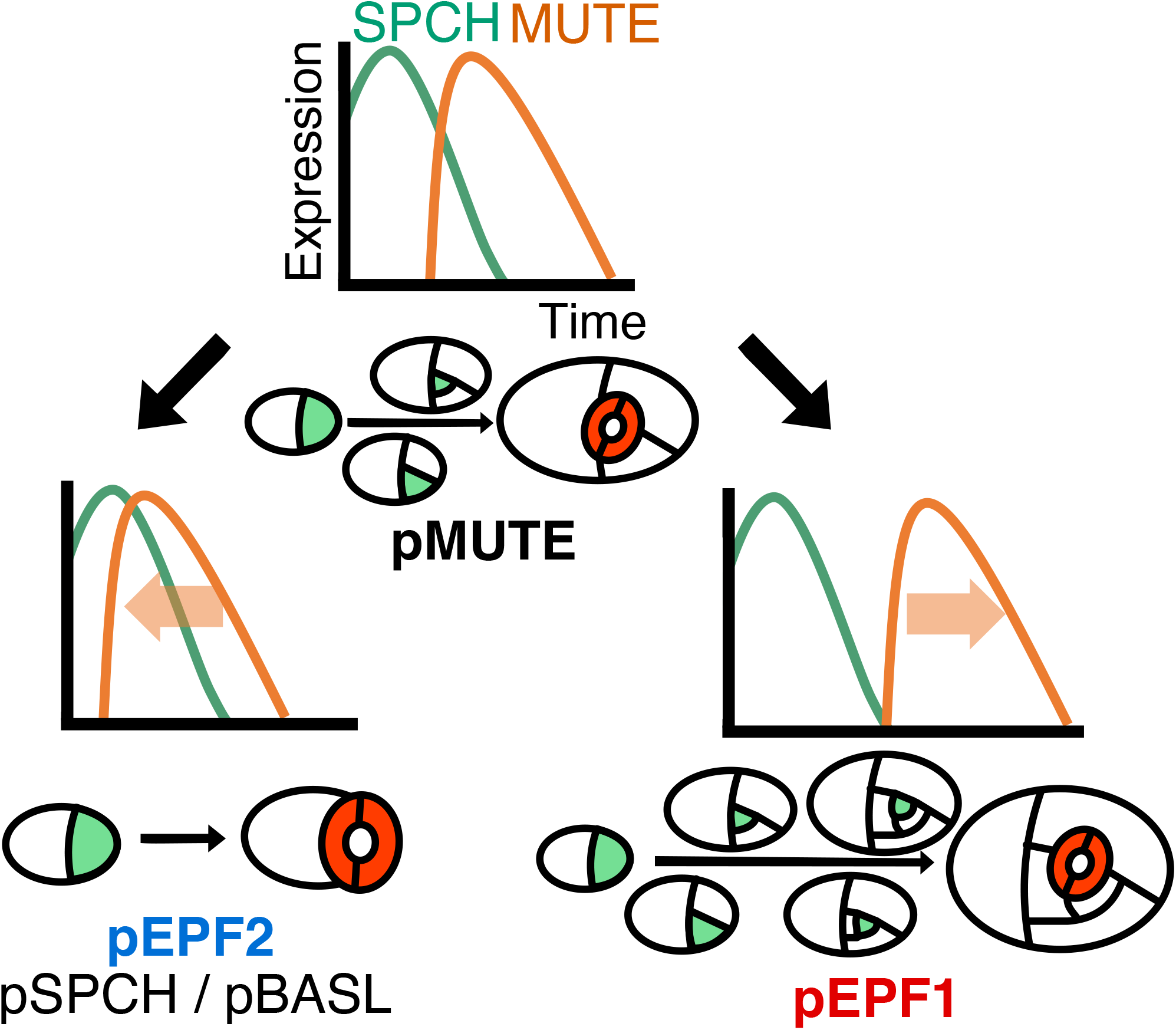
Graphical summary of the result. By modifying the temporal expression patter of MUTE, we successfully obtained the predicted behaviors of meristemoids in each line. Stomatal precursor cells are indicated in green, and GCs are indicated in red.

One of the notable findings of the present study is that delaying MUTE expression induced helicocytic stomatal development. Although there is no clear definition of helicocytic stomata, they are also found in some Brassicaceae species, such as the radish *Raphanus sativus* (Pant and Kidwai, 1967; Payne, 1970; Rudall et al., 2017). Thus, our model could be applied in future studies for comparing Arabidopsis and helicocytic Brassicaceae species in order to understand the origin of helicocytic stomatal development. Another notable result of this study is that non-contiguous stomatal cluster-like structures were frequently found in pEPF1 (Fig. 4C, H). The non-contiguous stomatal cluster is a specialized structure found in some helicocytic species, such as those in the genus *Begonia* (Rudall et al., 2017), and it has been considered to contribute to effective gas exchange and high water-use efficiency under certain environmental conditions (Papanatsiou et al., 2017; Tsai et al., 2022). Because there are also helicocytic species that do not have non-contiguous stomatal clusters, not only prolonged amplifying divisions but also some other factors are needed for the acquisition of the clustering phenotype. For example, a recent study suggested the possible involvement of TMM orthologs in the formation of the non-contiguous stomatal cluster in *Begonia*; this indicates the importance of altered spacing regulation in stomatal lineage cells (Tsai et al., 2022). The pEPF1 line established in this study would serve as excellent material for future evolutionary, physiological, and developmental studies on the formation of helicocytic stomata and non-contiguous stomatal clusters. For instance, it may be possible to infer the direct mechanisms behind the evolution of such traits by manipulating other stomata-related genes in the pEPF1 line and examining the resulting phenotypes.

Experiments in the three early MUTE expression lines produced in this study revealed some unexpected phenotypes such as excess cell divisions and aberrant spacing of stomata (Figs. 4−6, S3). The contiguous cluster of stomata observed in the early MUTE lines were in violation of the one-cell-spacing rule—the well-supported empirical rule in stomatal development (Sachs, 1991) (Figs. 4A, E, S4). In principle, this rule is only violated in mutants associated with developmental signaling (Berger and Altmann, 2000; Geisler et al., 2000; Bergmann et al., 2004; Shpak et al., 2005; Hara et al., 2007; Lampard et al., 2008; Dong et al., 2009) or under specific highly artificial environmental conditions (Akita et al., 2013). Therefore, the clustering stomata in pEPF2, pSPCH, and pBASL could be interpreted as a developmental defect in stomatal spacing that is probably related to the clustering MUTE-GFP expression observed in these lines (Figs. 3A, D, S3). We note here that a recent paper from a different research context reported the Arabidopsis line in which MUTE is driven by the *EPF2* promoter and reported that the line also formed contiguous stomatal clusters in its cotyledons (Smit et al., 2023), as in our pEPF2 line. The defective phenotypes of the early MUTE expression lines might be the result of technical limitations of the experimental system in this study. The three genes whose promoters were used here for inducing MUTE expression earlier are either critical regulators of the early patterning of stomatal lineage identity (*EPF2*, *BASL*) or the identity factor itself (*SPCH*). Therefore, driving MUTE by such promoters may interfere with the endogenous regulation network of stomatal patterning in Arabidopsis and produce unexpected phenotypes. Such a conflict in developmental signaling may also explain why many meristemoids continued to undergo devision in pEPF2 at 12 DAS (Fig. 6B), despite the increase in the number of cells expressing the division terminator MUTE (Fig. 4A, D, G). Alternatively, these defective spacing phenotypes may reflect an evolutionary constraint in Brassicaceae, in which meristemoid amplifying divisions are highly conserved (Pant and Kidwai, 1967). One possible mechanism for such a constraint might be the SLGC-mediated pathway for stomatal differentiation. In Arabidopsis, SLGC may regain the MMC fate to produce a new meristemoid through a process called spacing division (Pillitteri and Dong, 2013; Vatén et al., 2018; Ho et al., 2021), which was not observed during our analysis of *Callitriche* species (Doll et al., 2021a). In this process, early expression of MUTE in MMC and early meristemoid stages may be associated with MUTE expression in SLGCs that are regaining the MMC fate and may eventually lead to MUTE expression in both the meristemoid and its sister SLGC, and, thus, the formation of abnormally clustering stomata. Since contiguous clusters of stomata impair gas exchange and water-use efficiency (Dow et al., 2014; Lehmann and Or, 2015), genotypes with such traits are thought to be under negative selective pressures. This leads to the question of how anomocytic species without developmental defects, as seen in nature, evolve. Future comparative studies in both anomocytic and anisocytic species are needed to answer this question. Overall, the results in the early MUTE expression lines not only support the importance of temporal gene expression changes in the diversification of stomatal development, but also imply the possible presence of constraints on pattern diversification and the need for identifying additional factors involved in the evolution of patterns found in nature.

Since patterns of stomatal development have independently diversified in each lineage of plants, the diversification may be explained by various molecular mechanisms. We manipulated MUTE expression in this study to verify the previous model, but diversification of the stomatal development process may also be influenced by changes in SPCH expression. Moreover, results obtained here do not exclude the possibility that mechanisms other than the temporal shift in SPCH or MUTE expression may have driven the diversification of stomatal development patterns in some land plant lineages. For example, differences in the activity or stability of the SPCH protein may give rise to differences in the numbers of meristemoid divisions. In addition, modification of other factors present downstream or upstream of the SMF transcription factors, such as PEAPOD, which is known to regulate the activity of meristemoid divisions (White, 2006, 2022; Gonzalez et al., 2015), may also contribute to the diversification of stomatal development. Further, as recently proposed (Gong et al., 2022), the possible influence of cell size threshold on amplifying division termination may also explain differences in meristemoid behaviors among related species.

Previous evolutionary developmental biology (evo-devo) studies on stomata at the molecular level were largely limited to a small number of distantly related model species, such as moss and grass species (Chater et al., 2017; McKown and Bergmann, 2020), and a few of them explored evolution at the species level. Therefore, these studies could not elucidate the direct cause for the diversification of stomatal development in each lineage of plants. Further, previous studies used only indirect approaches to studying the evolution of diverse meristemoid behaviors, such as genomic comparison (Xu et al., 2018) and analyses of gene expression patterns (Doll et al., 2021a). The current study overcomes these limitations of previous studies by experimentally reproducing the possible evolutionary process and providing direct evidence for the molecular mechanisms related to pattern diversification. Since we successfully validated the model established in *Callitriche* (Plantaginaceae, asterids) (Doll et al., 2021a), which is distantly related to Arabidopsis (Brassicaceae, rosids), we believe that the model is applicable to broad lineages of plants. Deep and wide conservation of SPCH and MUTE in the genomes of land plants further supports this view (MacAlister and Bergmann, 2011; Ran et al., 2013). Thus, the model we experimentally validated here would provide a new framework for stomata evo-devo at the species level, and the findings of this study will serve as an important step toward understanding the independent evolutionary processes of stomatal development in various lineages of land plants.

## Abbreviations

GCs: guard cells
GMCs: guard mother cells
MMCs: meristemoid mother cells
DAS: days after sowing
SI: stomata index
SLGCs: stomatal lineage ground cells
PCI: precursor cell index

## Acknowledgments

The authors are grateful to Drs. Ayami Nakagawa and Keiko Torii (Nagoya University) for providing the materials, and Dr. Gorou Horiguchi (Rikkyo University) for assistance with the transformation of Arabidopsis.

## Author contributions

Y.D ., H.K., and H.T. designed the research. Y.D. performed the research and analyzed the data. Y.D., H.K., and H.T. wrote the paper.

## Funding

This work was supported by a Sasakawa Scientific Research Grant to YD (grant no. 2019-4112; the Japan Science Society), a Grant-in-Aid for Japan Society for the Promotion of Science (JSPS) Fellows to YD (grant no. JP20J20446; JSPS), a Grant-in-Aid for Early-Career Scientists to HK (grant no. JP20K15816; JSPS), and a Grant-in-Aid for Scientific Research on Innovation Areas to HT (grant no. JP19H05672; MEXT).

### Data availability

All the data supporting the findings of this study are included in the manuscript and supplementary figures.

